# Motor Learning Outside the Body: Broad Skill Generalisation with an Extra Robotic Limb

**DOI:** 10.1101/2025.06.27.661550

**Authors:** Maria Molina-Sanchez, Lucy Dowdall, Giulia Dominijanni, Viktorija Pavalkyte, Clara Gallay, Edmund da Silva, Dani Clode, Tamar R. Makin

## Abstract

Our ability to transfer motor skills across tools and contexts is what makes modern technology usable. The success of motor augmentation devices, such as supernumerary robotic limbs, hinges on users’ capacity for generalised motor performance.

We trained participants over seven days to use an extra robotic thumb (Third Thumb, Dani Clode Design), worn on the right hand and controlled via the toes. We tested whether motor learning was confined to the specific tasks and body parts involved in controlling and interacting with the Third Thumb, or whether it could generalise beyond them.

Participants showed broad skill generalisation across tasks, body postures, and even when either the Third Thumb or the controller was reassigned to a different body part, suggesting the development of abstract, body-independent motor representations.

Training also reduced cognitive demands and increased the sense of agency over the device. However, participants still preferred using their biological hand over the Third Thumb when given the option, suggesting that factors beyond motor skill generalisation, cognitive effort, and embodiment must be addressed to support the real-world adoption of such technologies.

## Introduction

In science fiction, robotic technology is often portrayed as a natural extension of the human body, with characters effortlessly adapting to new robotic limbs and intuitively controlling them across a wide range of scenarios. Such depictions suggest that artificial limbs can be seamlessly incorporated into human motor plans and generalised across diverse environments. In reality, the process is far more complex. Today, supernumerary robotic limbs, such as robotic fingers^1–3^, arms^4,5^, legs^6,7^, and tails^8^, are being developed to increase the body’s degrees of freedom while working in coordination with our biological limbs. While these devices perform well in controlled settings for specific tasks, their effectiveness in broader, real-world contexts remains uncertain^9,10^. This raises a fundamental question: how do we transition from today’s constrained, task-specific control to a future where artificial limbs are used as flexibly as our own biological ones? Realising this future will require a deeper understanding of the mechanisms that support, or hinder, the generalisation of motor skills with these technologies.

One possibility is that motor learning with extra robotic limbs primarily depends on developing somatosensory and motor skills in the body parts responsible for their control and interaction. As this form of learning is grounded in sensorimotor mechanisms, it may place fewer demands on cognitive resources and therefore be more intuitive. However, motor skills developed with the body are often inflexible and highly context dependent. For example, motor learning does not always transfer between hands^11–13^ or between orthogonal muscle groups^14^. Similarly, the transfer of tactile learning between fingers is often restricted to topographic neighbours or homologs^15,16^. Thus, if motor augmentation learning follows a similar pattern, it is likely that its transferability, and hence benefit, will be limited.

Alternatively, phenomena such as motor equivalence suggest that some motor tasks rely primarily on cognitive mechanisms, making them more adaptable. A well-documented example is handwriting, which shows a consistent motor output across different effectors (hand, foot or mouth), indicating an effector-independent motor representation^17,18^. Similarly, learning to use augmentation devices may involve developing a higher-level, abstract representation of movement, not tied to specific muscle groups or sensory inputs. Such learning could promote greater flexibility, allowing motor skills to generalise beyond body-specific motor and somatosensory constraints. However, it may also place greater demands on cognitive resources, which may limit the accessibility and usefulness of these technologies.

In this study, we aimed to explore whether motor skill acquired with an extra robotic thumb, the Third Thumb (Dani Clode Design), can generalise beyond the specific contexts in which it is acquired. We tested whether learning is confined to the specific tasks and body parts involved in controlling and interacting with the device during training, or whether skill learning affords generalisation beyond training parameters. The Third Thumb is worn on the right hand (effector) and controlled by pressure sensors activated by the left and right big toes (controllers), proportionally mediating two degrees of freedom, flexion and adduction. Using this device requires coordinating sensorimotor resources between the right hand and the toes; while motor commands are sent to the toes, sensory feedback about the outcome of these actions is primarily received from the right hand. Additionally, the motor plans of the right fingers have to be updated to integrate and coordinate effectively with the movement of the Third Thumb. This distinct right hand–toes sensorimotor loop offers a unique opportunity to dissect the relative contributions of the controllers and the effector to the generalisation of motor learning.

To assess the generalisation of motor skill, and the roles of controller and effector mappings in this process, we trained participants to use the Third Thumb over seven consecutive days. Training involved a diverse range of tasks designed to promote motor exploration and generalisation to novel scenarios. A control group of participants, who did not receive any training with the Third Thumb, was included for comparison. Before and after training, we conducted a battery of behavioural tests varying in task, body posture, and sensorimotor demands. We hypothesised that if motor learning relied on specific motor or somatosensory resources, trained participants would struggle to generalise their newly acquired skill when these demands were altered. Instead, our findings revealed that, following training, Third Thumb motor skill generalised across tasks, body postures, and sensorimotor mappings.

Furthermore, Third Thumb control became less cognitively demanding and was associated with an increased sense of agency over the device. Yet, participants still preferred using their biological hand over the Third Thumb when given the option, suggesting that factors beyond motor skill generalisation, cognitive effort, and embodiment must be addressed to support the real-world adoption of these technologies.

## Results

### Daily at-home training with the Third Thumb improves motor performance

We first assessed motor skill development with the Third Thumb in trained participants (augmentation group; N = 30) through a battery of motor tasks during 2-hour training sessions, conducted over seven consecutive days.

Since the effectiveness of these technologies relies on their usability outside the laboratory, we aimed to show that participants could effectively learn to use the Third Thumb at home with minimal supervision. Initial and final sessions (Day 1 and Day 7) were held in the lab, while the intermediate sessions (Day 2 to Day 6) were conducted remotely at participants’ homes, allowing participants some flexibility in when and how they practised (see Fig. 1 for experimental design). This approach enabled us to evaluate skill acquisition in less controlled, more naturalistic environments.

**Fig. 1.**
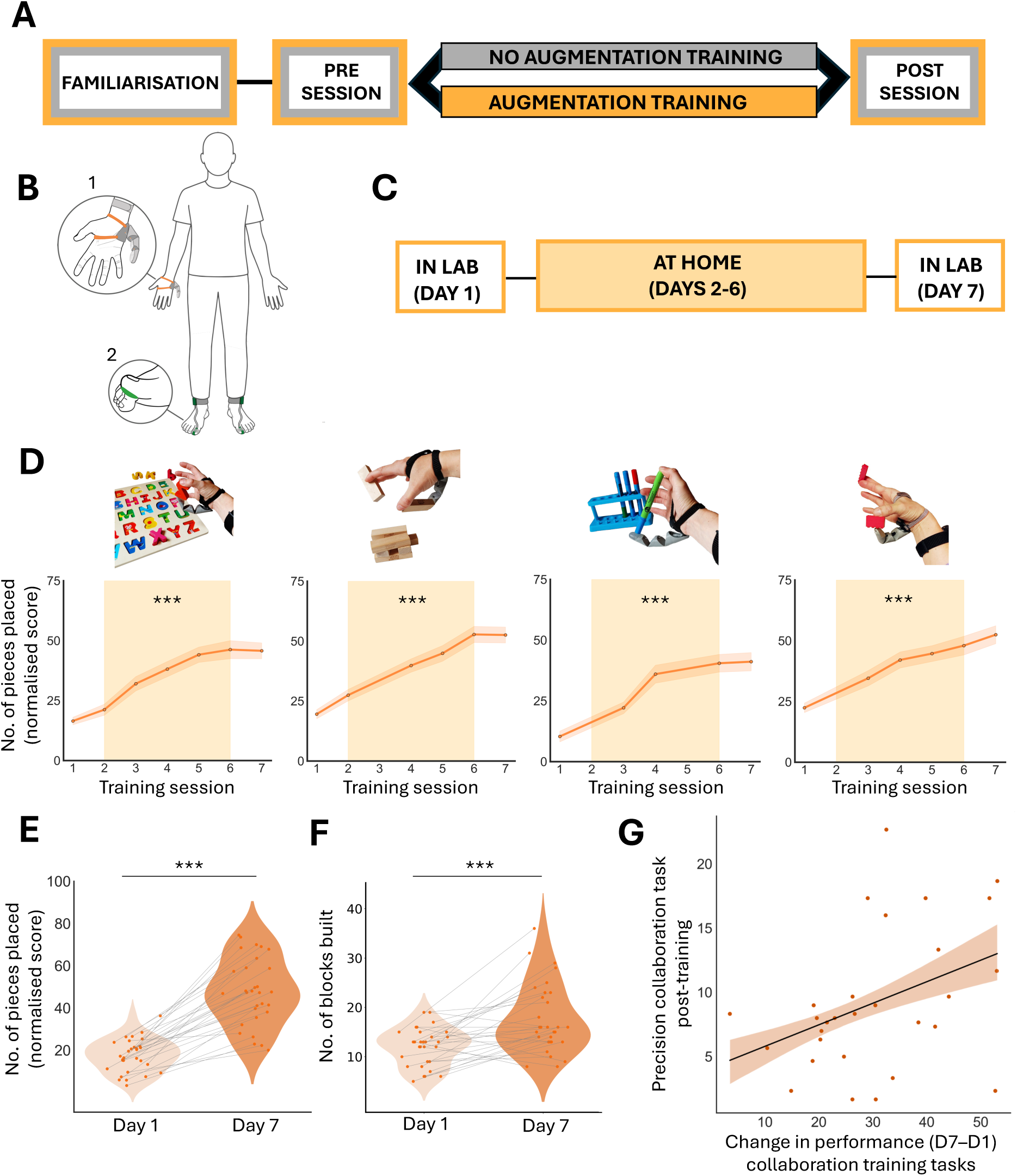
Training sessions for augmentation participants **(A)** Experimental design. All participants first completed a short familiarisation session with the Third Thumb, followed by a set of behavioural tests (pre-training session). They were then assigned to either the augmentation (N = 30) or control group (N = 40). Participants in the augmentation group, but not the control group, subsequently completed seven days of Third Thumb training. At the end of this period, all participants completed a post-training session, during which the same set of behavioural tests was repeated as in the pre-training session. **(B)** The Third Thumb is a 3D-printed robotic thumb, placed on the ulnar side of the right hand, actuated by two motors on a wrist strap (1). It is controlled by two pressure sensors underneath the big toes (2), allowing for proportionate control of flexion/adduction. **(C)** In-lab and at-home training timeline for augmentation participants. **(D)** Collaboration training tasks. Augmentation participants significantly improved on all collaboration tasks across sessions. Dots represent normalised group means. Faded orange around the line represents SEM. Background orange represents at-home training, where performance was self-reported by participants. Asterisks denote a significant effect of session (p < 0.001). **(E)** Augmentation participants significantly improved in the collaboration training tasks (scores combined in a single averaged value) from Day 1 to Day 7 (lab scores). Violin plots show the distribution of participant scores, with lines between dots reflecting within subject changes from Day 1 to Day 7. **(F)** Learning validation task. Augmentation participants improved their performance in an untrained strategy-free collaboration task between early (Day 1) and late (Day 7) stages of training. Asterisks denote main effect of day (p < 0.001). **(G)** Augmentation participants showed a significant correlation between their improvement on the training collaboration tasks (combined) between Day 1 and Day 7 and their performance in the precision collaboration task at the post-training session (standard configuration). The shaded orange area around the regression line represents SEM.

The training sessions included a diverse range of tasks, aimed at enhancing motor control with the Third Thumb and promoting generalisation through motor exploration. An overview of the training tasks, along with all other experimental tasks, their timeline, purpose, and sample sizes is provided in the Supplementary Material Table S1.

A key focus was on collaboration tasks, where the Third Thumb was used alongside a biological finger to grasp and transport items of various shapes and sizes to a set location (Fig. 1D). Participants repeated each collaboration task a minimum of five times across the seven sessions. The outcome measure was the average number of pieces successfully placed in one minute across trials, with scores being normalised for each task and participant. Highly significant improvements were observed in daily training sessions across all tasks (main effect of session for each task, *F* ≥ 29.827, *p* ≤ 0.001, *η*p² ≥ 0.611) and when comparing in-lab performance between Day 1 and Day 7 for each task (*t* ≥ 8.301, *p* < 0.001, *d* ≥ 1.569; Fig. 1D-E). For further comparisons, we combined the scores of these collaboration tasks to a single averaged value.

Additionally, participants were trained on other types of tasks, aimed at exploring different functional uses of the Third Thumb, such as independent gripping (individuation task), stabilising objects to increase the fingers manipulation capacity (split control task) and extending the hand’s gripping capacity (expansion task) (see Supplementary Fig. S1).

To validate the learning acquired during training, augmentation participants were assessed in a strategy-free task in which they were free to use the Third Thumb in any way they chose, either independently, or in collaboration with the augmented hand. The task involved constructing a tower from building blocks based on a template. Consistent with training gains, we observed a highly significant improvement in performance from Day 1 to Day 7 (*t*(29) = 4.42, *p* < 0.001, *d* = 0.808; Fig. 1F). Overall, these results highlight the efficacy of our largely home-based and minimally supervised training.

### Motor learning with the Third Thumb generalises to new task demands and body postures

Is motor learning with the Third Thumb confined to the specific tasks encountered during training? We introduced a hand–Thumb coordination task during early (Day 1) and late (Day 7) stages of training, to test whether participants could transfer their newly acquired motor skill to untrained tasks with new demands. Unlike the training tasks, which all involved object manipulation, this task required augmentation participants to perform a coordinated movement to bring their biological fingers into contact with the tip of the Third Thumb (Fig. 2A). This task was repeated twice: (i) seated, with full vision and (ii) seated with vision occluded. The outcome measure was the average number of successful hits between the instructed finger and the Third Thumb across trials. Participants demonstrated a significant improvement for both task variations from Day 1 to Day 7 (main effect of day, *F*(1, 25) = 8.752, *p* = 0.007, *η*p² = 0.259) with no significant interaction between day and task variation, *F*(1, 25) = 1.107, *p* = 0.303, BF10 = 0.420; Fig. 2B). The improvement under the vision-occluded condition suggests that training may enhance motor integration of Third Thumb movements with the hand, aligning with previous findings^1^.

**Fig. 2.**
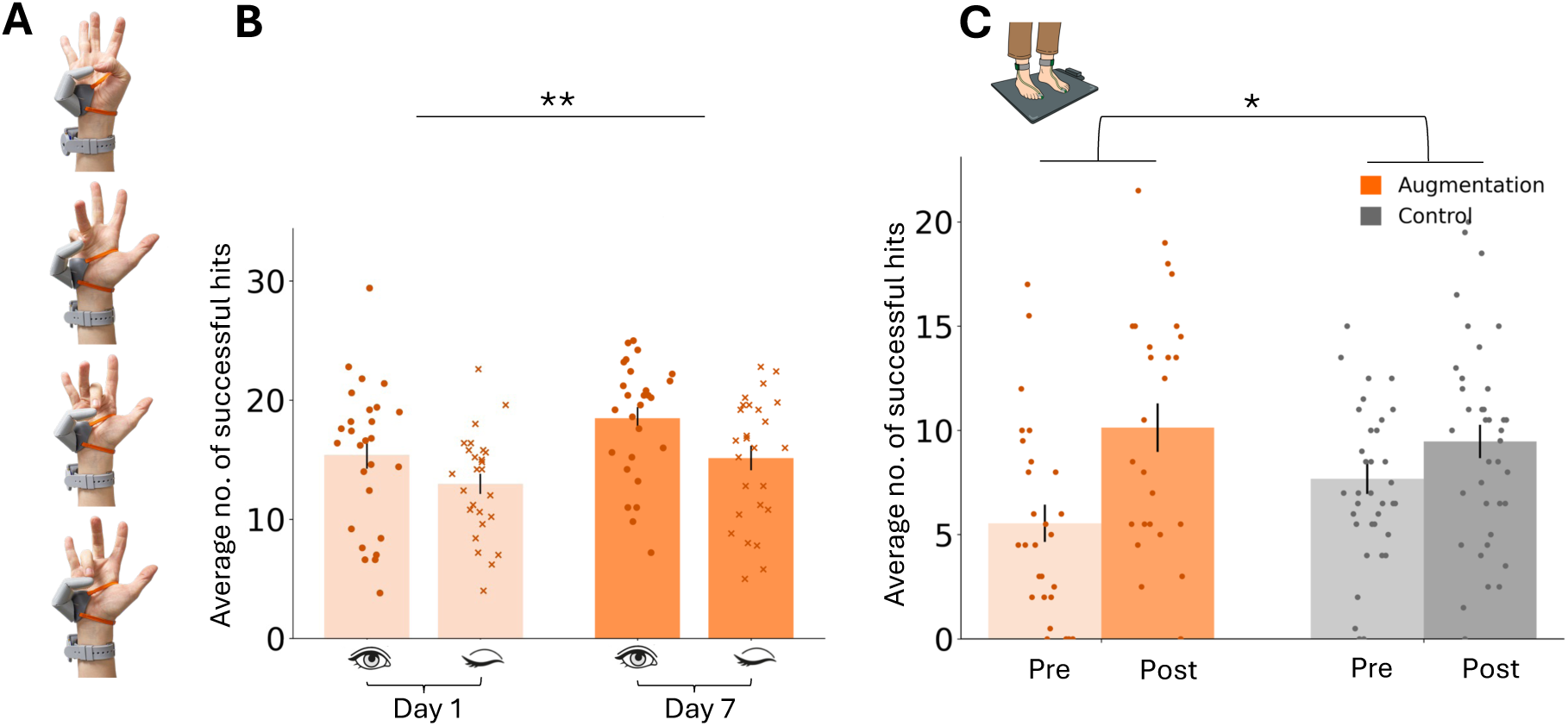
Generalisation to new task demands and postures **(A)** Hand–Thumb coordination task: participants used the tip of the Third Thumb to touch the tip of a randomly cued biological finger. **(B)** Augmentation participants improved their performance in the hand–Thumb coordination task between early (Day 1) and late (Day 7) stages of training, with eyes open (circles) and with vision occluded (crosses). Asterisks denote main effect of day (p < 0.01). **(C)** Augmentation (orange) and control participants (grey) improved their performance in the same coordination task from pre- to post-training sessions, performed while balancing; however, the augmentation group showed significantly greater gains. Asterisks denote an interaction effect (p < 0.05). Bars represent group means, error bars indicate SEM, and individual dots represent individual participants.

To further test the flexibility of Third Thumb motor skill, we asked whether participants could still effectively operate the device under increased postural demands. For this purpose, we introduced a task that required Third Thumb control while balancing on a balance board.

Since the device is operated via the toes, maintaining balance while using the Third Thumb places additional demands on the controller. To investigate this, a subset of augmentation (N = 26) and control participants (N = 38) were tested on the hand–Thumb coordination task (with full vision) while balancing on a balance board, during the pre- and post-training sessions. Here, we focus on performance in the hand–Thumb coordination task independently of balance ability, which is addressed separately below. Although both groups improved their performance across sessions (main effect of session, *F*(1, 57) = 26.614, *p* < 0.001, *η*p² = 0.318), a significant session x group interaction was observed, *F*(1, 57) = 5.671, *p* = 0.021, *η*p² = 0.090. This was driven by a greater improvement in the augmentation group (*t*(23) = 4.516, *p* < 0.001, *d* = 0.922) relative to the controls (*t*(34) = 2.319, *p* = 0.027, *d* = 0.392, Fig. 2D). To account for lower baseline performance by the augmentation group at the pre-training session, this analysis was repeated with baseline performance as a covariate, resulting in a similar trend (*F*(1, 56) = 3.452, *p* = 0.068, *η*p² = 0.058).

Together, these findings demonstrate that augmentation participants could successfully transfer their Third Thumb motor skill beyond the specific tasks and postures encountered during training.

### Motor learning with the Third Thumb generalises to new controller and effector mappings

Is motor learning tied to the specific body parts used for controlling (toes) and interacting (right hand) with the Third Thumb? We address this question by reassigning the roles of the controllers and the effector to a different body part. During pre- and post-training sessions, all participants completed a precision collaboration task, requiring them to pinch-grip small discs of paper between the Third Thumb and their biological thumb and stack them onto poles (Fig. 3A and Supplementary Video 1). The task was performed in three counterbalanced configurations: (i) standard — Third Thumb on the right hand, controlled by the toes (as used during the familiarisation session and training); (ii) heels — Third Thumb on the right hand, controlled by the heels; and (iii) left-hand — Third Thumb on the left hand, controlled by the toes. Performance was measured as the average number of discs stacked per minute, compared across sessions and against all control participants, to account for potential effects of task repetition.

**Fig. 3.**
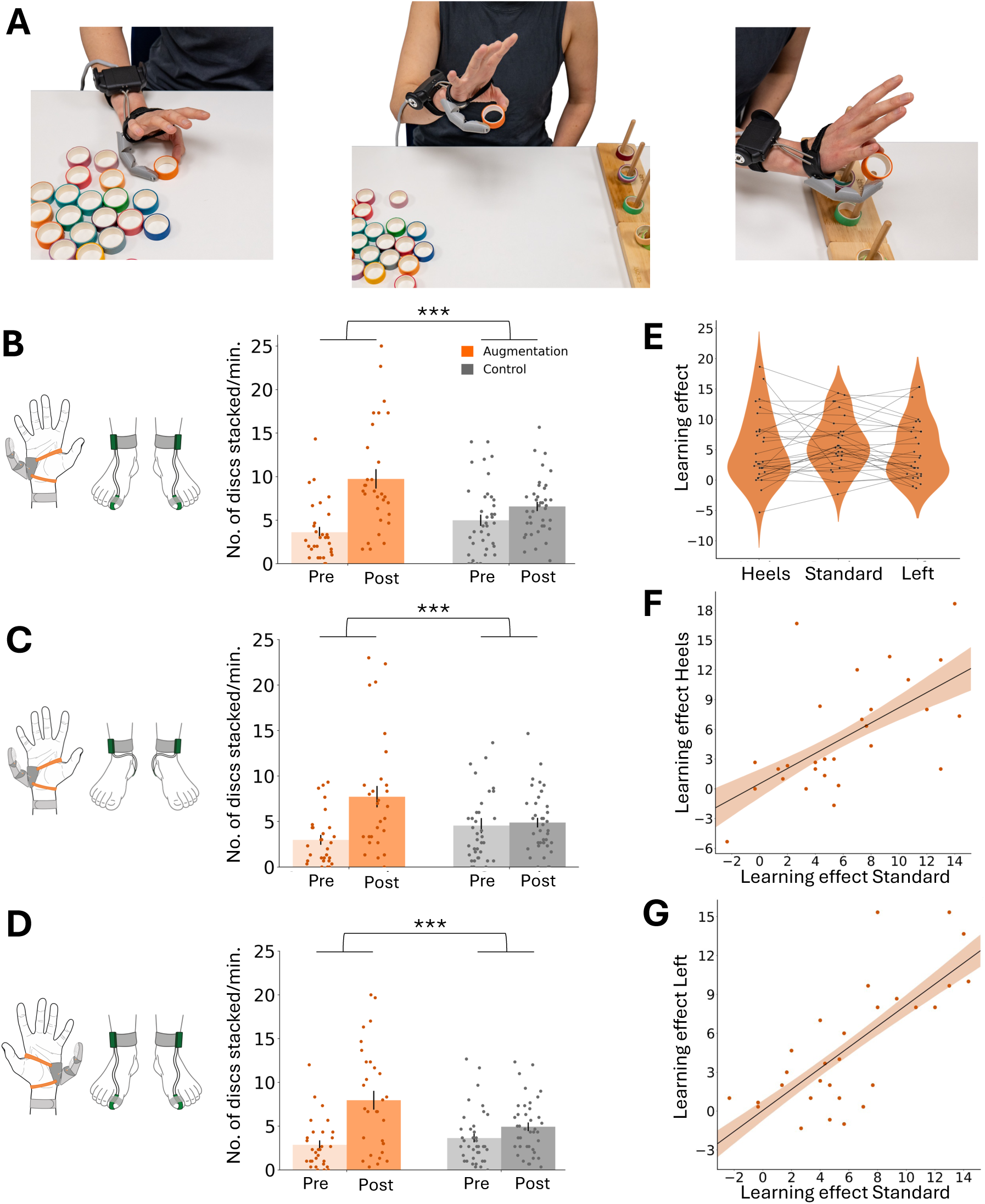
Generalisation to new sensorimotor configurations **(A)** Precision collaboration task (not included in the training set), where participants stacked small paper discs onto a pole. **(B)** Augmentation participants (orange) showed improved performance in this task compared to controls (grey) in the standard configuration (Third Thumb on the right hand, controlled by the toes), as well as **(C)** when using a new controller (heels configuration), and **(D)** a new effector (left-hand configuration). Bars represent group means; error bars indicate SEM. Asterisks denote significant interaction effects (p < 0.001). **(E)** Learning effects across the three configurations did not significantly differ within the augmentation group. Violin plots show the distribution of participant scores, with lines between dots reflecting within subject changes across the three configurations. **(F-G)** The learning effect with a new controller and a new effector strongly correlated with the learning effect in the standard configuration. For all sub-figures, individual dots correspond to participants’ scores. The shaded orange area around the regression lines represents SEM.

We found a significant main effect of session across all three configurations (*F* ≥ 28.940, *p* < 0.001, *η*p² ≥ 0.305), indicating that repetition improved performance. Crucially, we found highly significant group-session interactions, not only for the standard configuration (*F*(1, 66) = 18.361, *p* < 0.001, *η*p² = 0.218; Fig. 3B), but also for the new controller (heels configuration: *F*(1, 66) = 13.946, *p* < 0.001, *η*p² = 0.174; Fig. 3C) and the new effector (left-hand configuration: *F*(1, 65) = 14.872, *p* < 0.001, *η*p² = 0.186; Fig. 3D). This was driven by significant group differences in the post-training session, with the augmentation group showing superior performance compared to controls in each of the three configurations (*t* ≥ 2.449, *p* ≤ 0.017, *d* ≥ 0.601), indicating that it was Third Thumb training that selectively improved performance.

We then compared learning effects, calculated as the difference between post- and pre-training performance, across the standard, heels, and left-hand configurations in the augmentation group. Surprisingly, there were no differences between configurations (*F*(2, 54) = 0.831, *p* = 0.44, BF₁₀ = 0.20; Fig. 3E), indicating that participants’ skill gains in the heels and left-hand configurations were comparable to those in the standard configuration. This was further supported by strong correlations between the learning effect in the standard configuration and both the heels (*r*(28) = 0.58, *p* = 0.001; Fig. 3F) and left-hand configurations (*r*(29) = 0.74, *p* < 0.001; Fig. 3G). These results demonstrate that although participants trained in a specific controller–effector configuration, they showed comparable learning gains in new controller and effector mappings. Additionally, performance gains on the training collaboration tasks from Day 1 to Day 7 significantly correlated with post-training performance in the standard configuration (*r*(29) = 0.39, *p* = 0.039; Fig. 1G), suggesting that overall skill acquisition predicted subsequent task success.

Altogether, these findings suggest that motor skill acquired with the Third Thumb is not constrained by the specific effector or controller used, pointing to the formation of high-level motor representations that generalise across body parts.

### Training reduces cognitive reliance

Given that generalisation of Third Thumb motor skill did not depend on low-level sensorimotor training parameters, we next assessed whether generalised device control relies more on high-level cognitive attentional resources. During the pre- and post-training sessions, all participants completed a dual motor–cognitive task to assess the impact of increased cognitive load on motor performance. This involved performing the precision collaboration task (i.e., stacking paper discs) in the standard configuration, while simultaneously engaging in an arithmetic task (Fig. 4A). To ensure active engagement, participants were instructed that maintaining accurate counting in the arithmetic task was their primary goal. Outcome measures included: (i) the average number of discs stacked across trials, and (ii) average counting accuracy across trials.

**Fig. 4.**
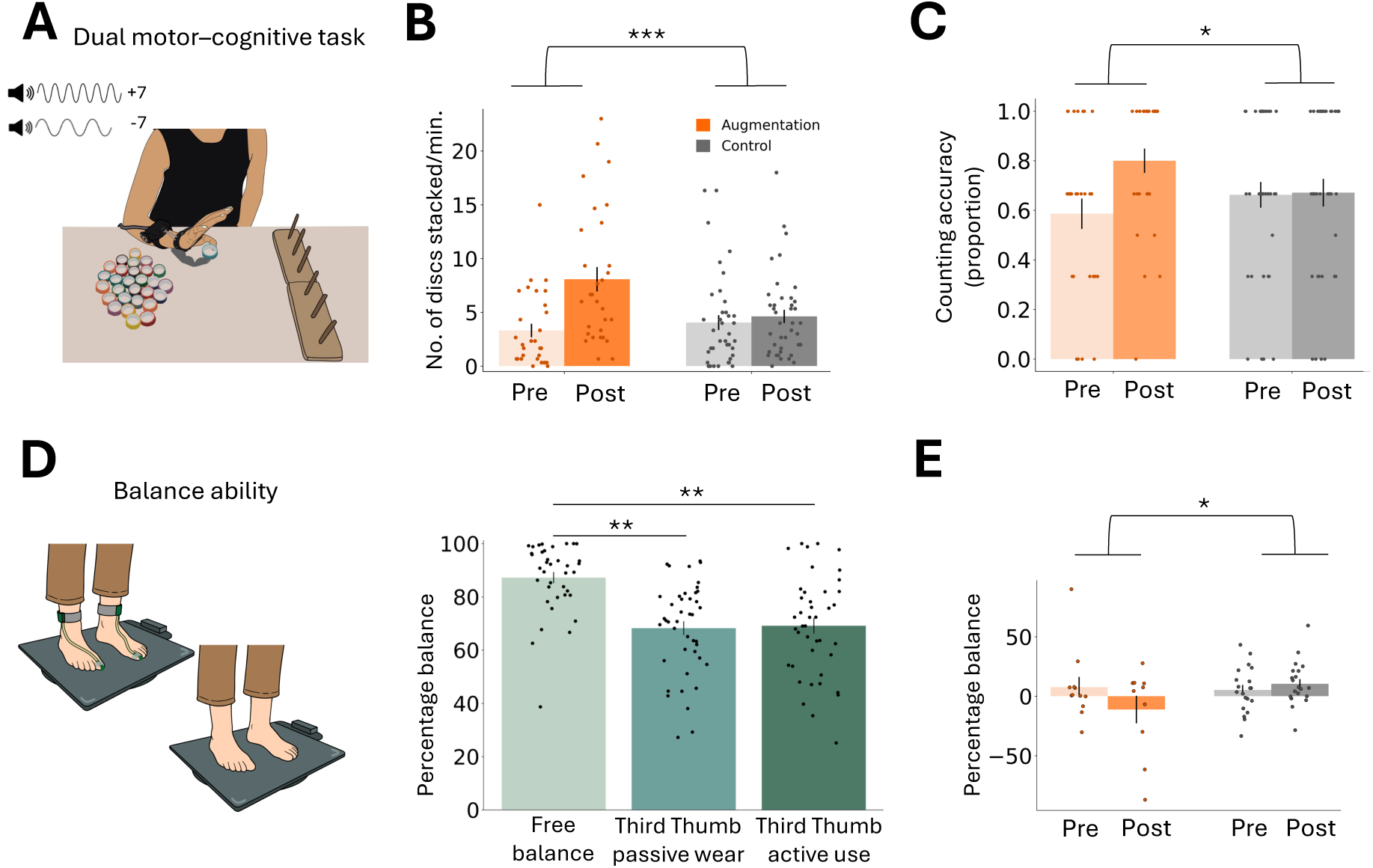
Third Thumb effect on cognitive reliance and lower limb function **(A)** Dual motor–cognitive task where participants completed the precision collaboration task (i.e., stacking paper discs onto poles) in the standard configuration, while simultaneously performing an arithmetic task aloud. **(B)** Augmentation participants showed improved motor performance in the dual task compared to controls. Asterisks denote a significant interaction effect (p < 0.001). **(C)** Augmentation participants demonstrated a greater improvement in the arithmetic task than controls, suggesting that training allowed them to allocate more cognitive resources to the arithmetic task. Asterisk indicates a significant interaction effect at p < 0.05. **(D)** Third Thumb’s impact on balance ability. When pooling all participants before training, we found a significant reduction in balance ability when either using (active use condition) or simply wearing the Third Thumb (passive wear condition), compared to free balance during familiarisation. Asterisks denote a significant effect of condition (p < 0.01). **(E)** Augmentation participants showed a significant decline in free balance following training compared to controls. Asterisk denotes significant interaction effect (p < 0.05). For all subfigures, bars represent group mean, error bars indicate SEM, and individual dots correspond to participants’ scores.

We first analysed participants’ motor performance in the dual task, regardless of their accuracy in the concurrent arithmetic task. Motor performance improved significantly across sessions (main effect of session, *F*(1,67) = 41.518, *p* < 0.001, *η*p² = 0.383; Fig. 4B).

Importantly, a highly significant group × session interaction (*F*(1, 67) = 18.806, *p* < 0.001, *η*p² = 0.219) was observed, driven by the augmentation group showing significantly better post-training performance relative to controls (*t*(67) = 2.772, *p* = 0.007, *d* = 0.673). This replicates motor gains due to Third Thumb skill learning in the precision collaboration task.

To determine the impact of cognitive load on motor performance, we compared participants’ motor performance in the dual task to the single motor task (i.e., without cognitive load).

Results revealed that participants stacked significantly fewer discs under cognitive load across sessions (main effect of cognitive load (*F*(1, 66) = 13.477, *p* < 0.001, *η*p² = 0.170). There was a significant cognitive load × session interaction (*F*(1, 66) = 4.041, *p* = 0.048, *η*p² = 0.058), driven by a greater impact of cognitive load on motor performance in the post-training (*t*(67) = –3.767, *p* < 0.001, *d* = 0.457, mean difference= –1.779) compared to the pre-training session (*t*(67) = –2.357, *p* = 0.021, *d* = 0.286, mean difference = –0.838). This may reflect a floor effect in pre-training motor performance, leaving limited scope for further performance reduction under cognitive load. There was no significant interaction between cognitive load and group (*F*(1, 66) = 0.678, *p* = 0.413, BF₁₀= 2.436×10⁻¹⁰), indicating that the arithmetic task similarly affected motor performance across both groups, despite the selective improvement in motor skill among augmentation participants.

We next considered performance on the cognitive component of the dual task. Arithmetic performance did not differ significantly between groups at either the pre-(*z* = –0.98, *p* = 0.325) or post-training session (*z* = 1.56, *p* = 0.124). However, the augmentation group showed significantly greater improvement than controls, *z* = 2.226*, p =* 0.027*, r* = 0.276; permutation-based Wilcoxon rank-sum test; Fig. 4C. These results suggest that, although training improved motor performance, it did not make participants less susceptible to increased cognitive load relative to controls. However, only augmentation participants showed improvements in arithmetic performance, indicating that training may have enabled a more efficient allocation of cognitive resources to the primary task.

### Wearing and controlling the Third Thumb may impact lower limb function

Given the broad generalisation observed, we asked whether using the toe-controlled Third Thumb could result in maladaptive transfer effects on lower-limb function. Specifically, learning to use the toes for Third Thumb control may negatively impact their original role in balance. We tested this hypothesis by measuring balance performance in a subset of participants (N = 47; augmentation: N = 16, controls: N = 31) across several conditions. Balance performance was quantified as the percentage of time the balance board was maintained within a tilt range of –5° to +5°.

To test whether using the Third Thumb, or simply wearing it, affects balance ability even before training, we pooled all participants and compared balance performance across three conditions: (i) active use — using the Third Thumb to perform the previously described hand–Thumb coordination task; (ii) passive wear — wearing the Third Thumb but not using it during the same task; and (iii) free balance — balancing without the device during the familiarisation session.

A significant main effect indicated differences in performance between the three conditions (*F*(2, 60) = 8.044, *p* < 0.001, *η*² = 0.211). As expected, we observed a modest but highly significant decline in balance ability while actively using the Third Thumb compared to free balance (*t*(31) = –3.225, *p* = 0.003, *d* = 0.570; Fig. 4D). Interestingly, merely wearing the Third Thumb without using it also led to a similar decline (*t*(32) = –3.109, *p* = 0.004, *d* = 0.541), with no significant difference found between the passive wear and active use conditions (*t*(41) = 0.077, *p* = 0.939, BF₁₀ = 0.167). This suggests that regardless of active use, the physical presence of the Third Thumb on the body may affect postural control. No group differences were observed before training (*F*(1, 30) = 0.274, *p* = 0.604, BF₁₀ = 0.388), and training did not improve performance when using or wearing the Third Thumb (non-significant session × group and condition × group interactions (*F*(1, 36) ≤ 0.529, *p* ≥ 0.472, BF₁₀ ≤ 0.182).

Next, to assess potential after-effects of Third Thumb training, we evaluated participants’ balance without the device (free balance) during the pre- and post-training sessions. Due to incomplete data caused by technical issues, a linear mixed model was employed, showing no significant main effect of session, *F*(1, 42.70) = 1.08, *p* = 0.304. However, a significant session × group interaction was observed, *F*(1, 42.70) = 5.33, *p* = 0.026. Follow-up simple effects contrasts showed no significant difference between groups at the pre-training session (*Z*(73.714) = 0.472, *p* = 0.639); however, following training, there was a significant decline for augmentation participants compared to controls (*Z*(73.725) = −2.644, *p* = 0.010). Note that with a repeated measures ANOVA, the session × group interaction was only a trend, *F*(1, 32) = 3.25, *p* = 0.081, η²ₚ = 0.092).

These findings suggest a potential mild after-effect on lower-limb function following extended use of the Third Thumb. However, given the small sample size, this result should be interpreted with caution.

### Training enhances a sense of agency over the Third Thumb

Does the development of flexible motor skill with the Third Thumb change participants’ sense of agency over the device? We explored this by analysing participants’ responses to four agency-related statements from an embodiment questionnaire, completed at the end of both the pre- and post-training sessions (for a list of the agency-related statements, see Supplementary Table S2; for additional embodiment questionnaire categories see ref.^19^. The outcome measure was the average rating across the four questions on a seven-point Likert-type scale. All participants reported a significant increase in their sense of agency across sessions (*F*(1, 59) = 54.825, *p* < 0.001, *η*p² = 0.482). Additionally, a significant group by session interaction was found, *F*(1, 59) = 4.595, *p* = 0.036, *η*p² = 0.072, driven by a higher increase in agency in the augmentation group (*t*(27) = 5.230, *p* < 0.001, *d* = 0.988, mean difference: 1.062), compared to controls (*t*(31) = 4.938, *p* < 0.001, *d* = 0.873, mean difference: 0.555; Fig. 5A).

**Fig. 5.**
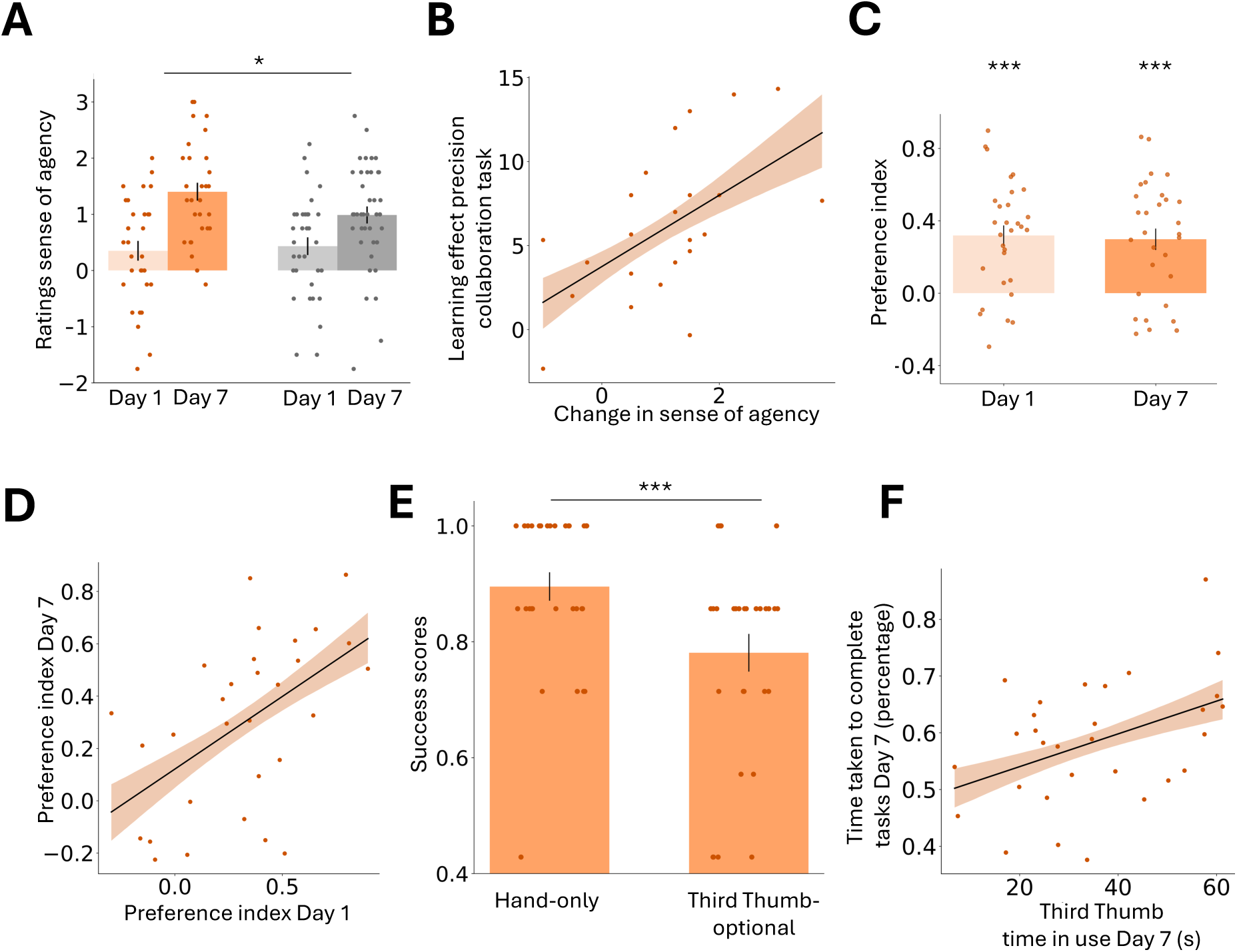
Third Thumb training effect in sense of agency and free-choice tasks **(A)** Augmentation participants showed a greater increase in sense of agency compared to controls. Asterisk denotes a significant interaction effect (p < 0.05). **(B)** In the augmentation group, increases in sense of agency across sessions correlated with skill gains in the precision collaboration task (standard configuration). **(C)** Positive preference index across sessions indicating a consistent preference for using the biological hand during free-choice tasks. Asterisks indicate a significant difference from zero (p < 0.001). **(D)** Preference indices were significantly correlated across sessions. **(E)** Participants were less successful in the free-choice tasks when they had the option to use the Third Thumb (Third Thumb-optional condition), compared to when they repeated the tasks using only their hand (Hand-only condition). Asterisks indicate a significant effect of condition (p < 0.001). **(F)** Greater use of the Third Thumb was associated with slower task completion.

We then examined whether the increase in sense of agency was related to the learning effect in the precision collaboration task (standard configuration) for augmentation participants. A significant correlation was found between the two (*r*(26) = 0.508, *p* = 0.008; Fig. 5B), suggesting that motor skill development is associated with increased sense of agency over the device.

### Skill acquisition did not change voluntary Third Thumb use in a set of free-choice tasks

Real-life multitasking often involves situations in which one hand is occupied while the other must perform tasks that typically require bimanual coordination (e.g., plugging in a USB connection, opening a jar). These contexts could, in principle, benefit from the support of an additional limb. Still, it remains unclear whether individuals will choose to apply their augmentation skills in such challenging situations, or whether they will continue to rely on their biological body alone. To investigate whether augmentation participants would choose to apply their generalised Third Thumb skill, we asked them to perform a series of everyday object manipulation challenges (e.g., unfolding a pair of socks, plugging in a USB connection; see Supplementary Video 2), in which they could in principle benefit from using the Third Thumb, although its use was not strictly required. Participants completed the tasks once during early (Day 1) and late (Day 7) training. Each task had a time limit for completion, and no specific instructions were given other than participants had to use their augmented (right) hand, with the option to incorporate the Third Thumb if they chose to (Third Thumb-optional condition). On Day 7, participants also completed the same task battery without the Third Thumb (Hand-only condition).

To quantify participants’ tendency to rely on either their biological hand or the Third Thumb during the Third Thumb-optional condition, we computed a preference index based on offline video analysis of task performance. Positive values indicate more time spent using the hand alone, while negative values reflect a preference for incorporating the Third Thumb to complete the task. Overall, participants preferred to adapt using their own hand rather than rely on newly acquired Third Thumb skill, as shown by a positive preference index on both Day 1 and Day 7, *t*(29) ≥ 5.027, *p* < 0.001, *d* ≥ 0.918; Fig. 5C, with no change across sessions, *t*(29) = 0.370, *p* = 0.714, BF₁₀ = 0.207. Preference indices were strongly correlated across these two sessions, *r*(29) = 0.526, *p* = 0.003; Fig. 5D, suggesting that inter-individual differences were driven by individuals’ baseline preference for using the Third Thumb.

Moreover, preference changes across sessions were not correlated with participants’ motor gains in the precision collaboration task, *ρ*(28) = 0.11, *p* = 0.594, BF₁₀= 0.48, indicating that skill acquisition alone was not sufficient to shift these baseline preferences.

Importantly, Third Thumb preference negatively aligned with task success, calculated as the average percentage of tasks completed within the time limit. On Day 7, participants were significantly less successful when given the option to use the Third Thumb compared to using only their hand, *W*(29) = 168.500, *p* < 0.001, *r* = 0.971; Fig. 5E. Moreover, greater reliance on the Third Thumb was associated with slower task completion, *r*(29) = 0.427, *p* = 0.019; Fig. 5F, suggesting a trade-off between incorporating the device and task efficiency. Participants appeared to respond to this trade-off rationally, opting for the strategy that allowed them to complete tasks more efficiently. This suggests that, under time pressure, users prioritised efficiency over novelty.

For reference, the same task was also completed by a subset of controls (N = 20). Although both groups improved their success rate across sessions (main effect of session, *F*(1,48) = 15.924, *p* < 0.001, *η*p² = 0.249), there was no session × group interaction, *F*(1,48) = 0.019, *p* = 0.89, BF₁₀ = 0.260, indicating that performance gains likely reflected task familiarity rather than training. Similar patterns were found in task completion time. Together, these findings suggest that, within the time scale tested, the Third Thumb does not generalise as effectively as the hand in adapting to motor challenges.

## Discussion

Much of the discourse in motor augmentation has focused on identifying the best body parts and related biosignals for wearing and controlling these devices^5,9,20–25^. This focus reflects classic motor control studies showing that learning is typically tied to the muscles, postures, and sensory conditions under which it occurs, with generalisation across effectors often limited^13,14,26,27^. However, our findings suggest that motor augmentation learning may not be as tightly constrained by anatomical location as traditionally assumed. Following training, participants generalised their Third Thumb skill across tasks, body postures, and even when either the effector or controller was remapped to a different body part, pointing to the development of a flexible, effector- and controller-independent internal model. This flexibility parallels the phenomenon of motor equivalence observed in complex, lifelong skills like handwriting, where consistent outcomes across different effectors suggest that movements are represented in the brain at an abstract level rather than as muscle-specific commands^17^. This is even more surprising, considering that aspects of this skill acquisition may have involved *de novo* learning processes, which require the construction of entirely new control policies not derived from prior motor experience^28,29^. That such generalisation emerged after only brief training in adulthood highlights the potential for rapid motor augmentation adaptation, with promising implications for real-world applications in rehabilitation, productivity enhancement, and assistive care.

The training paradigm was intentionally designed to support generalisation by fostering self-directed motor exploration across varied task demands. Participants were exposed to a broad repertoire of tasks intended to reflect the different motor challenges they might encounter in daily life. These tasks varied in strategy, the degree of collaboration with the biological fingers, the properties of objects used, and the specific ways participants interacted with them. Importantly, while each task had a clearly defined goal, participants received minimal instructions on how to achieve it, and most of the training took place at home without direct supervision. This encouraged exploration of movement strategies and allowed some flexibility in when and how they practised. Research shows that diverse training conditions enhance generalisation by promoting flexible, transferable learning rather than rigid, context-specific responses^30,31^. In parallel, motor learning studies have shown that granting learners greater autonomy, such as through self-controlled practice, can boost motivation and engagement, leading to deeper and more effective learning^32^. Altogether, this suggests that it is not motor augmentation itself that diverges from typical motor learning, but rather the flexible, and arguably more ecological, training approach (relative to classical lab-based motor learning tasks) that facilitated the extensive generalisation observed.

The broad generalisation observed raised concerns about potential maladaptive transfer effects. Given the toe-control Third Thumb’s mechanism, we assessed whether its use might interfere with lower-limb function via a balance task. We found reduced balance performance both when the Third Thumb was actively used and when it was merely worn. However, we do not interpret this as a maladaptive consequence of motor learning generalisation, as the effect was already present prior to training. Rather, it suggests that simply wearing an augmentation device can broadly influence motor behaviour, likely due to implicit adjustments arising from the dual role of the toes in both device control and postural stability^33^. This aligns with research showing that the motor system can adapt to overlapping or secondary demands, even when these are not the direct focus of a task, leading to unintended behavioural changes^34^. In our case, participants may have unconsciously altered their toe posture to minimise accidental activation of the device, thereby compromising balance. Nonetheless, a possible indication of maladaptive consequences from skill learning with the Third Thumb comes from a trend towards a session by group interaction in the free balance task, driven by a decline in balance performance across sessions in the augmentation group relative to controls. Although this finding should be interpreted cautiously, it raises the possibility that motor adaptations developed during augmentation might persist beyond the period the device is worn. This highlights the need for further research into the long-term effects of motor augmentation on broader motor functions^35,36^.

Although we hypothesised that generalisation across sensorimotor mappings would engage high-level cognitive mechanisms, findings from the cognitive load task reveal a more nuanced picture. While motor performance was affected by cognitive load, this effect was not specific to training, as both groups were similarly impacted. However, we observed an improvement in the cognitive component of the dual task among Third Thumb participants, suggesting that training facilitated more efficient allocation of cognitive resources. This does not contradict our hypothesis that cognitive processes are involved early in learning, but rather indicates that, with practice, device control can become more automatic. This aligns with evidence that motor skills requiring substantial cognitive engagement, such as those acquired through explicit processes, can become automatised over time^37^. Similar patterns have been reported in prosthesis studies, where arbitrary control strategies, initially more cognitively demanding than biomimetic ones, led to reduced cognitive load following training^38^.

Despite substantial learning, broad generalisation, and increased embodiment, participants did not preferentially choose to use the Third Thumb when given the option during free-choice tasks. One likely explanation is that they opted for the most efficient strategy, favouring their biological hand, which allowed for faster and more reliable task completion compared to the newly acquired skill with the Third Thumb. The time-constrained nature of the tasks, designed to reflect real-world conditions where time pressure often favours efficiency, may have further influenced this choice. Although participants demonstrated significant skill acquisition, seven days of training have likely not been sufficient for the Third Thumb to surpass the motor capability of the biological hand. However, the finding that post-training preference to use the Third Thumb significantly correlated with pre-training preference, rather than with motor skill gains, suggests that efficiency was not the sole factor influencing participants’ choices. Whether extended use might shift these preferences, leading to greater adoption of augmentation devices in daily life, is yet to be determined.

While augmentation devices may not offer immediate advantages for general, time-constrained daily activities, where the body, especially with two functional hands, remains highly efficient, they may still hold significant value in more complex or demanding contexts. Tasks requiring simultaneous manipulation and stabilisation, or multi-directional coordination, could particularly benefit from supernumerary limbs, especially given our findings that cognitively demanding tasks can be performed concurrently with Third Thumb use.

Additionally, individuals with motor impairments who are unable to use prostheses, such as those with hemiparesis following stroke^39–42^, spinal cord injury^43,44^, or other conditions affecting the function of an existing limb^45^, may achieve enhanced functional capacity through augmentation. Beyond practical applications, these technologies may also serve roles in self-expression, creativity, or aesthetic exploration^46^. In such contexts, adoption may be driven less by efficiency and more by personal identity, social trends or curiosity. This underscores the broader societal relevance of motor augmentation and suggests that factors beyond performance optimisation may increasingly shape personal preferences for these technologies.

In conclusion, our findings suggest that the key bottlenecks for the realisation of motor augmentation are not skill acquisition or training, but rather individual preferences, highlighting the critical role of human factors in the development of future technologies for the augmented body.

## Materials and Methods

### Participants

A total of 70 healthy participants (42 females; mean age = 25.04 ± 4.35; all right-handed) were recruited for this study from internet-based advertisements on the MRC Cognition and Brain Sciences Unit and the University of Cambridge SONA participant pools. Participants were randomly assigned to either the augmentation (N = 30; 19 females, mean age = 24.7 ± 3.91) or the control group (N = 40, 23 females, mean age = 25.3 ± 4.68).

All volunteers had no recognised motor or neurological disorders. Professional musicians and individuals who had more than five years of musical experience or had acquired musical training within the last five years were excluded from the study. Ethical approval was granted by the Cambridge Psychology Research Ethics Committee (PRE.2022.068) and the University College London Research Ethics Committee (REC: 12921/001). All participants gave their written informed consent before participating in the study.

An additional 20 participants were recruited but were excluded from the study due to withdrawal, incomplete datasets, or excessively long intervals between their pre- and post-training sessions (see *osf.io/c76xd*).

### Third Thumb description and mechanism of action

The Third Thumb (Dani Clode Design, Cambridge, UK) is a 3D-printed robotic thumb designed to enhance the motor capacities of the hand. Positioned on the ulnar side of the right hand, opposite to the biological thumb, it is actuated by two motors on a wrist strap (Fig. 1B). The Third Thumb allows for two independent movements: flexion-extension and adduction-abduction, controlled via pressure sensors placed underneath the user’s big toes. Pressure from the right toe induces flexion across the hand, while pressure from the left toe enables adduction toward the fingers, with the extent of movement proportional to the applied pressure. Two versions of the Third Thumb were used in this study. The *wired* version connects the motors and pressure sensor controllers via wires to a microcontroller and is powered by a wall plug, as detailed elsewhere^47^. This version was used for all seated tasks during the pre- and post-training sessions. The *wireless* version transmits signals from the toe pressure sensors to the motors via a microcontroller worn on the upper arm^1^ (Fig. 1B). Battery-powered, it provides users with greater flexibility to operate the device in unstructured environments, including unsupervised sessions outside the lab. This version was used for training and the balancing task during the pre- and post-training sessions. Both versions were worn and operated in the same manner.

### Experimental design

We used a longitudinal study design spanning 10 to 12 days (see *osf<u>.io/c76xd</u>* for experimental detail). All participants underwent the following in-lab sessions: (i) a 1.5-hour familiarisation session with the Third Thumb, which included an introduction to the equipment and the behavioural tasks, as well as a baseline assessment of their balance ability; and (ii) two behavioural testing sessions, each lasting 2.5 hours, with one at the beginning (pre-training) and another at the end of the experiment (post-training; Fig. 1A). The average interval between the pre- and post-training sessions was 10.77 ± 11.73 days across all participants, with no differences between the augmentation and control groups (*t*(68) = 1.201, *p* = 0.234, BF10 = 0.450). Additionally, participants in the augmentation group underwent seven daily 2-hour training sessions over consecutive days with the Third Thumb, scheduled between the pre- and post-training sessions. The training sessions in Day 1 and Day 7 were conducted in the lab, while sessions from Day 2 to Day 6 were conducted remotely at home with minimal supervision (Fig. 1C). The Day 1 training session could take place up to seven days after the pre-training session, while the Day 7 was conducted either the day before or on the same day as the post-training session. Beyond the structured daily sessions, augmentation participants were encouraged to use the Third Thumb as much as possible outside of training hours.

While participants in the control group did not receive training with the Third Thumb, half of them (N = 20) were provided with a foldable piano keyboard and underwent seven consecutive days of 2-hour keyboard training sessions, equivalent to the Third Thumb training regime. This training was facilitated using the interactive Yousician application (version 1.1, Yousician Ltd, Helsinki, Finland; see *osf<u>.io/c76xd</u>* for details). During the training sessions, participants learned to use the keyboard with their right hand while synchronising their keystrokes with toe presses on a connected pedal. This engaged them in hand-toe coordination movements like those performed by the augmentation group (see ref.^19^ for further details and training outcomes). The other half of the control group participants received no training. As no significant differences in performance were observed between these two subgroups (see *osf<u>.io/c76xd</u>*) their scores were combined into a single control group.

Throughout all sessions, Third Thumb tasks were performed exclusively with the augmented (right) hand. With the exception of one pre- post-training task conducted while balancing on a balance board, all other tasks were carried out while seated at a desk, with participants’ feet placed on a platform and their left hand resting on their lap.

### Augmentation group training sessions

Participants in the augmentation group underwent training on a set of motor tasks designed to explore a range of movements possible with the Third Thumb and to promote generalisation of motor skill. A strong emphasis was placed on collaboration tasks, where the Third Thumb worked in conjunction with one of the biological fingers to grasp objects of various shapes and sizes. There were four collaboration tasks in total, which included completing an alphabet puzzle, building a tower with wooden blocks, stacking markers, and sorting plastic building blocks (Figure 1D). The training also included an individuation task, where the Third Thumb interacted independently with a relevant object (e.g., hooking tapes) while the hand handled other task-irrelevant objects. Additionally, there was an expansion task, where the Third Thumb was used to extend the hand’s gripping ability (e.g., moving multiple objects), and a split control task, in which the Third Thumb was used to stabilise objects, thereby freeing the biological fingers to extend the hand’s manipulation capacities (see the Supplementary Fig. S1).

For each task, participants were seated in front of a computer displaying interactive videos via Edpuzzle (basic version, Edpuzzle Inc. Barcelona, Spain). This platform provided task instructions and allowed participants to self-record their scores. The minimally supervised training sessions conducted at home were recorded using Zoom (version 5.14.11, Zoom Video Communications, San Jose, California) and were occasionally monitored live. Each training session was structured to not exceed 2 hours, with individual tasks performed for a duration of 5 to 10 minutes and revisited across five to seven distinct training days.

### Training tasks scores analysis

To assess the progress of participants in the augmentation group for each training task, we normalised their scores using the min-max normalisation method. We then calculated the average score for each participant and each training session across tasks. Given the slight variations in the training regimen among participants, particularly regarding the distribution of tasks across different days, we organised the average scores according to the sequence of task repetitions. This approach involved categorising the scores based on whether it was the first, second, third, etc., repetition of the task, regardless of the specific days on which these repetitions occurred.

### Daily challenges and additional practice beyond structured training

At the end of each training session, augmentation participants completed a daily challenge, a short and engaging ecological task that incorporated some of the movements practised during training. These daily challenges aimed to encourage participants to continue using the Third Thumb beyond structured training by integrating it into realistic, everyday activities. Six different daily challenges were used throughout the training sessions, including swiping book pages, dealing cards, building a puzzle, hanging key rings, playing a dice game and imitating gestures. Each challenge was presented at the end of a session and tested in the subsequent session, allowing participants the opportunity to practice it outside of structured training hours.

Additionally, participants were encouraged to explore new ways to integrate the Third Thumb into their daily routines and to invent new tasks for its use. They self-reported their Third Thumb usage outside of structured sessions daily, including wear duration and examples of activities. The average usage duration for each participant was 5.23 ± 5.30 hours across the five days of at-home training.

### Learning validation task

At the end of the in-lab training sessions on Day 1 and Day 7, participants completed a strategy-free task aimed to validate the learning acquired with the Third Thumb. This task was not further practised but was used to assess early and late performance during training. Participants were required to build a tower following a provided template, using the given building blocks. They had 5 minutes to complete the task using the Third Thumb to carry the building blocks in any manner they preferred, either independently or in collaboration with the fingers of the augmented hand. The task was performed only once per session. The outcome measure was the number of building blocks correctly placed, regardless of whether the colours matched those in the original template.

### Testing protocol for generalisation of learning

To investigate skill generalisation, we employed a battery of behavioural tests to compare performance either between early (Day 1) and late (Day 7) training stages or between the pre- and post-training sessions. We evaluated the generalisation of Third Thumb skill to untrained scenarios using (1) new task demands (hand–Thumb coordination), and (2) a precision collaboration task with modified sensorimotor configurations.

#### (1) Hand–Thumb coordination task

To assess participants’ ability to generalise their newly acquired Third Thumb skill, we introduced a hand–Thumb coordination task that was not included in the training. In this task, augmentation participants used the Third Thumb to touch the tip of a randomly selected finger on their augmented hand (Fig. 2A). A MATLAB script generated a random sequence of target fingers, communicated through auditory cues. The experimenter recorded each successful hit between the Third Thumb and the cued finger, which triggered the cue for the next target. The task consisted of five one-minute runs and was performed on Day 1 and Day 7, first with full vision, then with vision occluded.

Additionally, participants performed a variation of this task while simultaneously balancing on a balance board during the pre- and post-training sessions. For this task, a MATLAB App generated a random sequence of target fingers, including a rest condition in which participants were instructed to maintain balance without attempting to perform the task. The sequence was presented via auditory cues: first indicating the target finger, followed by a “go” cue. Participants had three seconds to complete the specified finger hit, after which a “stop” cue signalled either successful completion or the end of the three-second period. This task was performed over two successive runs.

For each variation (with full vision, vision occluded, and while balancing), we calculated the average number of successful finger hits across runs.

#### (2) Precision collaboration task

To investigate the key elements of motor skill learning with the Third Thumb, we used an untrained precision collaboration task during the pre- and post-training sessions (see Supplementary video 1). Participants created a pinch grip between the Third Thumb and their biological thumb to pick up and stack small discs of paper onto poles. Sixty discs were placed to the participants’ right, with two three-pole wooden stackers fixed to their left, spaced 35 cm apart. Each participant completed three 1-minute runs, and performance was measured as the average number of tapes successfully stacked per run.

This task was performed in three different configurations, in a counterbalanced order: (i) in the standard configuration, participants wore the Third Thumb on the right hand and controlled it with their toes (thus, replicating the familiarisation and training conditions). (ii) In the heels configuration, participants controlled the Third Thumb using their heels instead of their toes. (iii) In the left-hand configuration, the Third Thumb was worn on the untrained (left) hand and controlled with the toes.

Finally, to assess the cognitive load associated with Third Thumb use, participants performed the precision collaboration task in the standard configuration while simultaneously completing an arithmetic task engaging their working memory, a method previously used to evaluate cognitive load in artificial limb control^1,38,48,49^. Participants began with a random number (81-180) and were then presented with a series of high- and low-pitch tones occurring randomly every 3-6 seconds. They were instructed to add 7 after each high-pitch tone and subtract 7 after each low-pitch tone, verbally reporting each result during a 1- minute run repeated three times. To ensure engagement in the cognitive task, participants were told that their primary goal was to maintain accurate counting. Outcome measures included: (i) the average number of tapes stacked across the three trials, and (ii) counting accuracy, assessed by whether their final count matched the correct result at the end of each trial.

The arithmetic task was initially practised independently (without the disc stacking task) during the familiarisation session to ensure participants could manage it. If participants failed the task in two 1-minute runs, the difficulty was adjusted from 7 to 3 or 1 for the subsequent pre- and post- training sessions, resulting in 59 participants using 7, 10 participants using 3 and 1 participant using 1.

### Impact on lower limb capacity

To explore potential functional costs of using the toe-controlled Third Thumb, we assessed its impact on balance performance in a subset of participants (N = 47). Balance was measured in three conditions: (1) while actively using the Third Thumb during the hand– Thumb coordination task in the pre- and post-training sessions, (2) during rest periods of the same task, when the device was worn but not used, and (3) during a 1-minute free-balance task without the device, conducted at familiarisation, pre-, and post-training sessions.

Balance performance was monitored using an inertial measurement unit (IMU; BWT61CL, WitMotion, China), with the primary outcome measure being the percentage of time participants maintained the balance board within an angle of −5 to +5 degrees.

### Embodiment and experience questionnaires

Participants’ subjective experience of agency over the Third Thumb was measured by examining their agreement with four agency-related statements drawn from an embodiment questionnaire, completed at the conclusion of both the pre- and post-training sessions.

Participants rated these statements on a 7-point Likert scale, ranging from −3 (strongly disagree) to +3 (strongly agree). The questionnaire also included statements addressing other explicit (phenomenological) aspects of embodiment, as detailed elsewhere^19^, along with four control (catch) statements, not expected to elicit positive responses regarding embodiment. Scores were calculated by averaging responses within the agency category for each participant. Three of the four agency statements were scored such that higher scores indicated an increased sense of agency; however, for one statement, lower scores reflected a stronger sense of agency (see a list of the statements in Supplementary Table S2).

Responses for that statement were reversed prior to averaging. If a participant’s average rating for the catch statements was above zero, their data for that session were excluded, resulting in the removal of one participant.

Participants also responded to three additional statements from an experience questionnaire: one assessed enjoyment of using the Third Thumb during the study, while the other two addressed potential use in daily life (see Supplementary Table S3). All responses were rated on the same 7-point Likert scale used in the embodiment questionnaire, with the potential use in daily life score calculated as the average of the two relevant statements. Results for these statements are reported in the supplementary material (Fig. S2).

### Free-choice tasks

To assess the extent to which participants would choose to apply their newly acquired Third Thumb skill, we designed a set of seven tasks involving the challenging manipulation of everyday objects. Although the use of the Third Thumb was not required to complete the tasks, it could enhance performance. The tasks included plugging in a USB connection, unfolding a pair of socks, opening a jar and pouring its contents, placing an elastic band around a jar, unzipping a zipped bag and retrieving its contents, moving five tennis balls, and flipping six upside-down wine glasses using as few movements as possible (see Supplementary Video 2). These tasks were performed by both the augmentation group and a subset of controls (N = 20) and were repeated once at the early (Day 1) and late (Day 7) stages of training. Participants were given a fixed time window to complete each task but were otherwise free to take as much time as needed within that limit. They were instructed to use only the augmented hand and could choose whether to use the Third Thumb (Third Thumb-optional condition). On Day 7, participants also completed the same battery of tasks without the Third Thumb (Hand-only condition).

Performance was recorded using three Logitech MSIP-REM-DZL-V-U0040 cameras positioned at different angles to provide a full view, with recordings managed through Open Broadcaster Software (OBS, version 29.1.3, Lain Bailey, Michigan, US). Performance analysis was based on live observation and video review. For the Third Thumb-optional condition, offline we measured the total time the Third Thumb and the hand were in use and calculated a preference index as follows: (hand-only – hand-Third Thumb)/ (hand-only + hand-Third Thumb). Additionally, for both the Third Thumb-optional and Hand-only conditions, we recorded whether each task was completed within the time limit (success rate), as well as the total time taken to complete it.

### Statistical analyses

All statistical analyses were performed using JASP (v.0.18.3). Normality was assessed using the Shapiro–Wilk test. When normality assumptions were met, parametric tests were used; otherwise, equivalent non-parametric tests were applied. Within-subject comparisons were conducted using paired-samples t-tests or two-tailed Wilcoxon signed-rank tests. Between-group comparisons used repeated-measures ANOVA with group as a fixed effect, and pre-training session score as a covariate when baseline differences were present. Where assumptions were violated, two-tailed Mann–Whitney tests or permutation-based Wilcoxon rank-sum tests were used. One-sample t-tests were used to assess whether values significantly differed from zero. For the free-balance task, due to missing data, a linear mixed model was employed with a random intercept for participant. Fixed effects included session and group. The model was estimated using Satterthwaite’s method with Type III sum of squares for p-value computation. All nonsignificant results were further evaluated using Bayesian tests (Cauchy prior width r = 0.707), with BF₁₀ < 1/3 interpreted as support for the null, BF₁₀ > 3 as support for the alternative, and values in between considered inconclusive^50^. Pearson or Spearman correlations were used depending on normality.

## Acknowledgements

We are grateful to all of our participants for their time and dedication. We thank Roderick Spender, Klara Selén, Katarina Krajnovic, Hannah Browning, Yuval Amichay, Karan Salvi, Ema Jugovic, Alexandra Williams and Mabel Ziman for their support with data collection and initial data pre-processing; Roni Maimon-Mor and Samuel McDougle for helpful comments on the initial manuscript; Silvestro Micera and Solaiman Shokur for helpful advice throughout the study; Yousician for providing us with a premium version of their application for this research. Funding: The study was funded by UKRI’s Frontier Research Guarantee (EP/X040372/1), the Engineering and Physical Sciences Research Council (EP/W004062/1), and the Medical Research Council (MC_uu_00030/10), awarded to T.R.M. T.R.M. was further partially funded by a Wellcome Trust Senior Research Fellowship (215575/Z/19/Z). G.D. was funded by European Union’s Horizon 2020 MSCA Programme under Grant Agreement (813713). Author Contributions: M.M. and T.R.M. led the conceptualisation and experimental design of the training paradigm and all behavioural tasks, except for the tasks involving balance performance. L.D. and G.D. supported the experimental design of the training paradigm. G.D. and T.R.M. led the conceptualisation and design of the tasks involving balance performance. D.C. designed and constructed the Third Thumb. M.M., L.D., G.D., V.P., and C.G. contributed to data collection for all datasets included in this manuscript. D.C. and E.dS. provided technical support during data collection. V.P. and C.G., supported by L.D. and M.M, contributed to the organisation and pre- processing of the training data and the generalisation tasks. M.M. and T.R.M. led the analysis of the training data and all behavioural datasets, except for the balance tasks, which were led by G.D. and T.R.M. M.M. also supported analysis of the balance tasks. M.M. and T.R.M. wrote the manuscript. M.M., supported by D.C., prepared the figures. L.D., G.D., V.P., C.G., E.dS., and D.C. provided feedback on the manuscript. T.R.M. provided funding for the work. Competing interests: The authors declare that they have no competing interests. Data and materials availability: The data that support the findings of this study will be available from the Open Science Framework upon publication (osf.io/c76xd). For the purpose of open access, the authors have applied a Creative Commons Attribution (CC BY) licence to any Author Accepted Manuscript version arising from this submission.

## Supplementary Materials

**Fig. S1.**
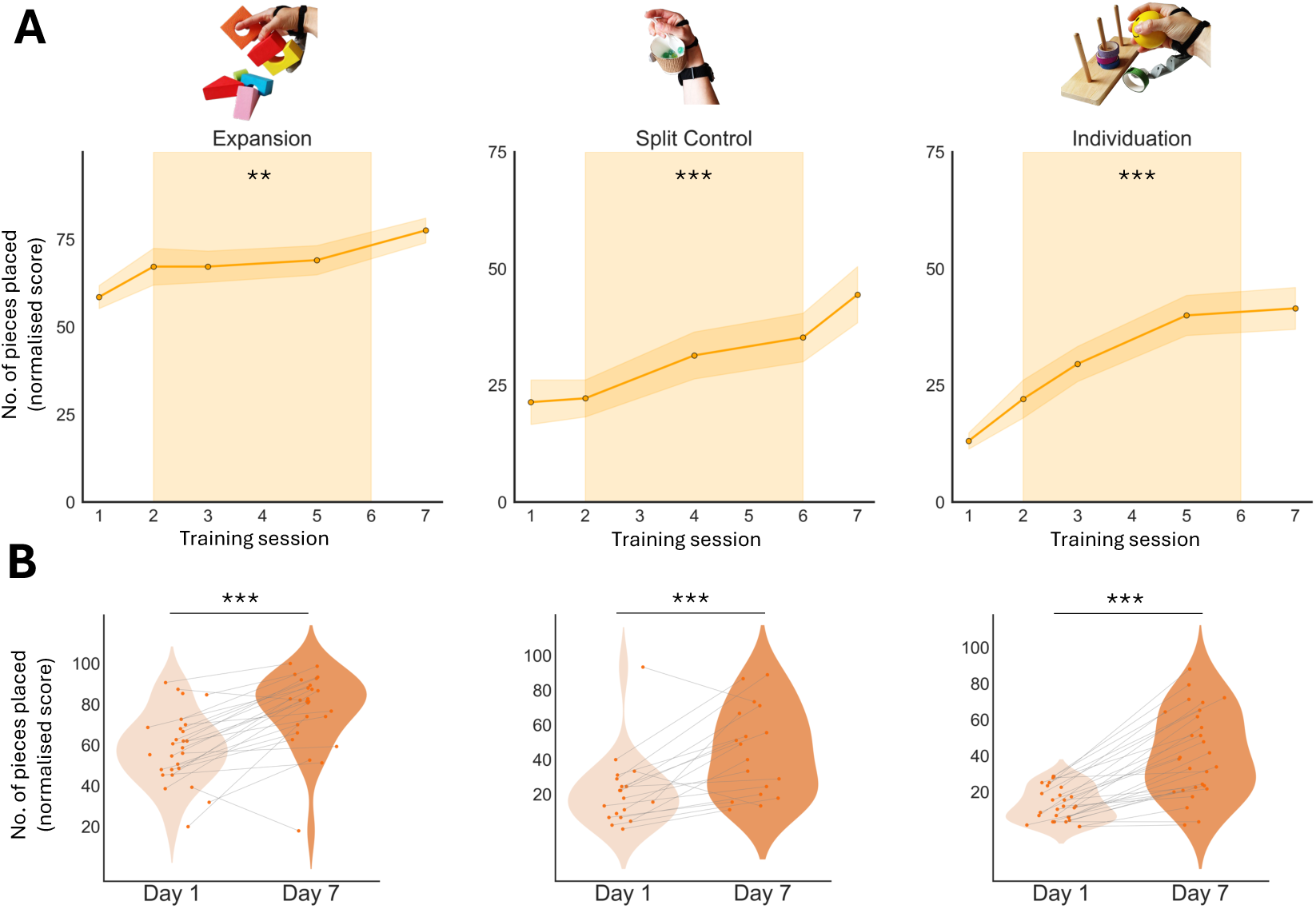
Motor performance across non-collaboration training tasks **(A)** Augmentation participants showed highly significant improvements over time in the Expansion, Split Control, and Individuation tasks (F ≥ 5.10, p ≤ 0.002, ηp² ≥ 0.35). Shaded orange areas around the lines indicate SEM. The light orange background represents the period of at-home training. Asterisks indicate a significant effect of session (**p < 0.01, ***p <0.001). **(B)** A significant increase was also observed when comparing in-lab performance during early (Day 1) and late (Day 7) stages of training for each task (p < 0.001). Violin plots show the distribution of participant scores, with lines between dots reflecting within subject changes from Day 1 to Day 7.

**Fig. S2.**
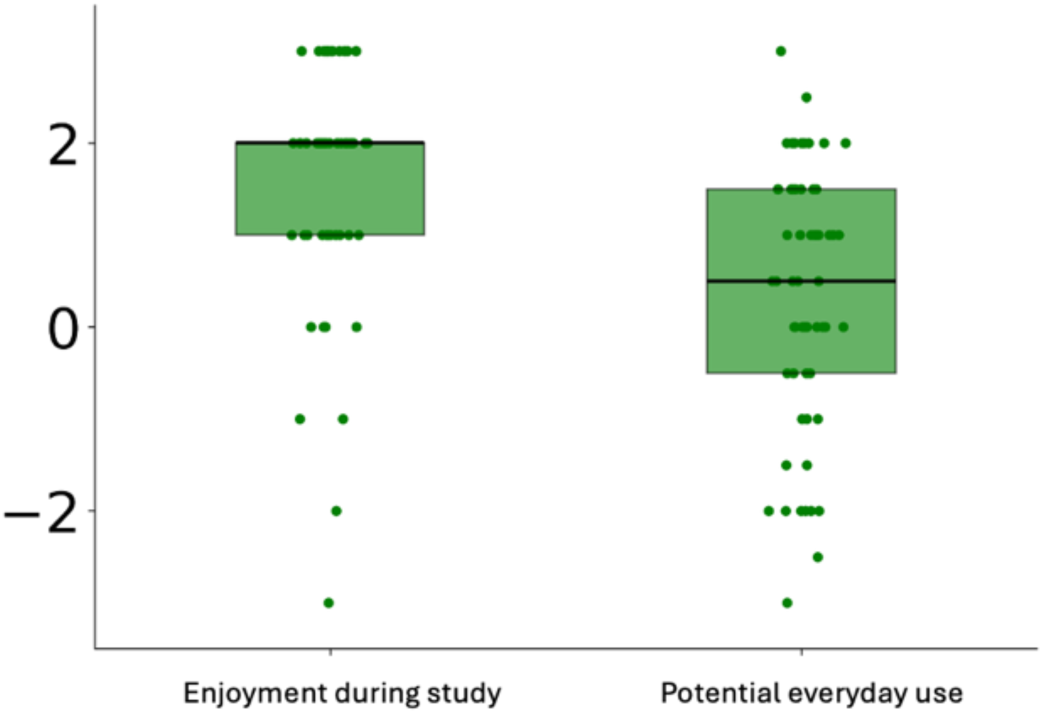
Experience questionnaire All participants reported a positive experience with the Third Thumb during the study, W(62) = 1617.500, p < 0.001, median = 2, mode = 2. However, they were more neutral about potential everyday use, W(62) = 907.0, p = 0.046, median = 0.5, mode = 0. We found no group differences in participants’ ratings for either category: enjoyment, U(61) = 521.0, p = 0.611, BF₁₀ = 0.260, and potential everyday use, U(61) = 460.0, p = 0.721, BF₁₀ = 0.289. Box plots represent the median and interquartile range (25th to 75th percentile). Individual dots correspond to participants’ scores. Asterisks indicate a significant difference from zero (p < 0.001, p < 0.05).

---

**Table S1.** Overview of experimental tasks, timeline, purpose, and sample sizes.

| Task | When | Purpose | Acquired sample | Additional Exclusions |
| --- | --- | --- | --- | --- |
| Collaboration training tasks:<br>- Alphabet puzzle<br>- Building a tower with wooden blocks<br>- Stacking markers<br>- Sorting plastic building blocks | Day 1 to Day 7 | Third Thumb skill learning | N=30 | N=1 due to missing data |
| Non-collaboration training tasks<br>- Individuation task<br>- Split control task<br>- Expansion task | Day 1 to Day 7 | Third Thumb skill learning | N=30 (Individuation)<br>N=19 (Split control)<br>N=28 (Expansion) | N=3 (Individuation) due to missing data<br>N=2 (Split control) due to missing data<br>N=4 (Expansion) due to missing data |
| Strategy-free task | Day 1 and Day 7 | Validation of learning | N=30 (Augmentation) | N=1 due to missing data |
| Hand–Thumb coordination task seated (eyes open and eyes closed) | Day 1 and Day 7 | Generalisation of learning | N=28 (Augmentation) | N=3 due to Third Thumb technical issues |
| Hand–Thumb coordination task while balancing | Pre- and post-training sessions | Generalisation of learning | N=26 (Augmentation)<br>N=38 (Control) | N=3 (Augmentation) and N=3 (Controls) due to IMU technical issues. |
| Precision collaboration task (Standard configuration) | Pre- and post-training sessions | Generalisation of learning | N=30 (Augmentation)<br>N=40 (Control) | N=1 (Augmentation) due to outlier performance and N=1 (Control) due to non-compliance with task instructions |
| Precision collaboration task (Heels configuration) | Pre- and post-training sessions | Generalisation of learning | N=30 (Augmentation)<br>N=40 (Control) | N=1 (Augmentation) because of incomplete data due to Third Thumb technical issues and N=1 (Control) due to non-compliance with task instructions |
| Precision collaboration task (Left-hand configuration) | Pre- and post-training sessions | Generalisation of learning | N=30 (Augmentation)<br>N=40 (Control) | N=1 (Augmentation) due to outlier performance and N=2 (Control) due to non-compliance with task instructions and Third Thumb technical issues |
| Dual motor-cognitive task (motor performance: tapes stacking) | Pre- and post-training sessions | Cognitive load assessment | N=30 (Augmentation)<br>N=40 (Control) | N=1 (Augmentation) due to Third Thumb technical issues.<br>N=1 (Control) due to non-compliance with task instructions |
| Dual motor-cognitive task (cognitive performance: arithmetic performance) | Pre- and post-training sessions | Cognitive load assessment | N=30 (Augmentation)<br>N=40 (Control) | N=1 (Augmentation) and N=4 (Controls) due to computer technical issues and missing data |
| Balance while wearing and using the Third Thumb | Pre- and post-training sessions | Impact on lower limb function | N=16 (Augmentation)<br>N=31 (Control) | N=3 (Augmentation) and N=4 (Control) due pressure mat/IMU technical issues |
| Free Balance | Familiarisation, pre- and post-training sessions | Impact on lower limb function | N=16 (Augmentation)<br>N= 31 (Control) | N=3 (Augmentation) and N= 7 (Control) due to pressure mat/IMU technical issues |
| Embodiment questionnaire | Pre- and post-training sessions | Sense of agency assessment | N=30 (Augmentation)<br>N=39 (Control) | N=1 (Augmentation) due to positive scores on catch trials.<br>N=7 (Control) due incomplete datasets |
| Experience questionnaire | Post-training session | Enjoyability and potential daily use | N=27 (Augmentation)<br>N=36 (Control) | N/A |
| Free-choice tasks<br>- Plugging in a USB connection<br>- Unfolding pair of socks<br>- Opening a jar and pouting its contents<br>- Placing an elastic band around a jar<br>- Unzipping a zipped bag and retrieving its contents<br>- Moving five tennis balls<br>- Flipping six upside-down wine glasses | Pre- and post-training sessions | Generalisation of learning | N=30 (Augmentation)<br>N=20 (Control) | N/A |

**Table S2.**
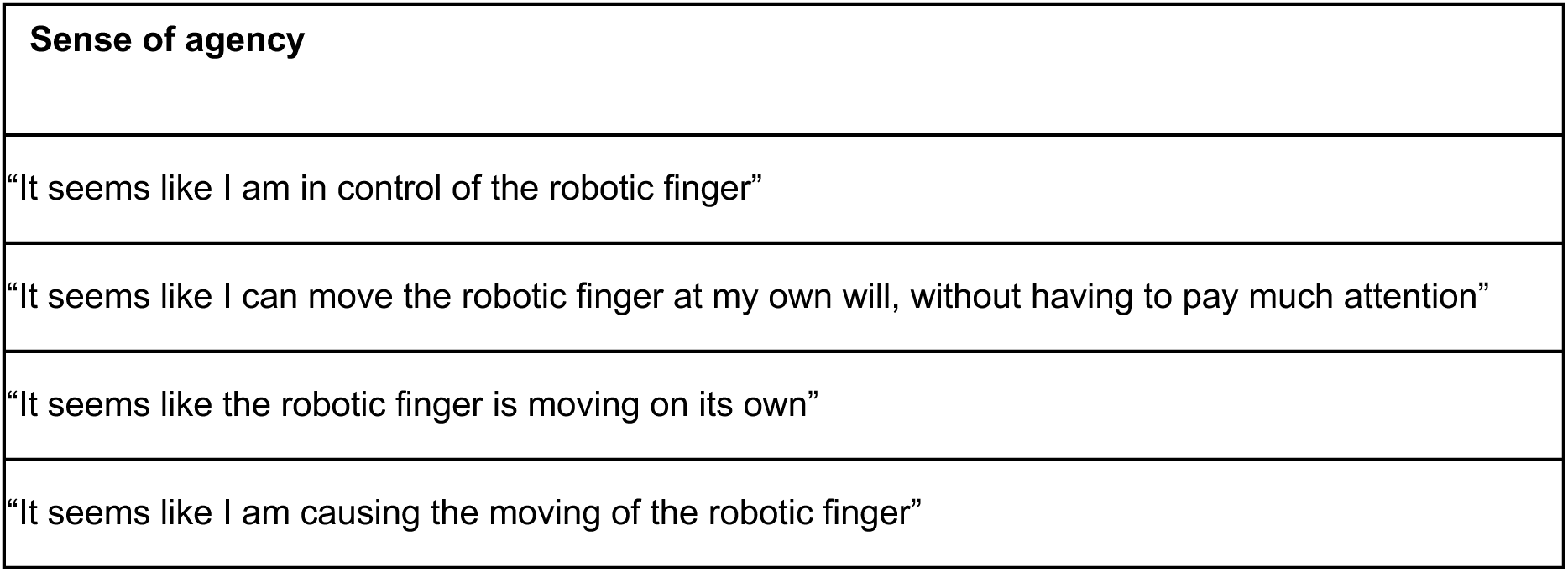
Sense of agency statements from the embodiment questionnaire.

**Table S3.** Enjoyability with the Third Thumb and potential everyday use statements from the experience questionnaire.

|  |
| --- |
| **Enjoyment during the study** |
| "I found the experience of having an extra robotic finger enjoyable" |
| **Potential use in daily life** |
| "I think having an extra robotic finger would be useful in my daily life" |
| "I think having an extra robotic finger would be enjoyable in my daily life" |

## References

1. Kieliba, P., Clode, D., Maimon-Mor, R. O. & Makin, T. R. Robotic hand augmentation drives changes in neural body representation. *Sci*. Robot. 6, eabd7935 (2021).

2. Shafti, A., Haar, S., Mio, R., Guilleminot, P. & Faisal, A. A. Playing the piano with a robotic third thumb: assessing constraints of human augmentation. Sci. Rep. 11, 21375 (2021).

3. Wu, F. & Asada, H. Supernumerary Robotic Fingers: An Alternative Upper-Limb Prosthesis. Proc. ASME Dyn. Syst. Control Conf. 2, V002T16A009 (2014).

4. Buratti, S. et al. Effect of Vibrotactile Feedback on the Control of the Interaction Force of a Supernumerary Robotic Arm. Machines 11, 1085 (2023).

5. Dominijanni, G. et al. Human motor augmentation with an extra robotic arm without functional interference. *Sci*. Robot. 8, eadh1438 (2023).

6. Hao, M., Zhang, J., Chen, K., Asada, H. & Fu, C. Supernumerary Robotic Limbs to Assist Human Walking With Load Carriage. J. Mech. Robot. 12, 061014 (2020).

7. Khazoom, C., Caillouette, P., Girard, A. & Plante, J.-S. A Supernumerary Robotic Leg Powered by Magnetorheological Actuators to Assist Human Locomotion. IEEE Robot. Autom. Lett. 5, 5143–5150 (2020).

8. Abeywardena, S., Anwar, E., Charles Miller, S. & Farkhatdinov, I. Mechanical Characterization of Supernumerary Robotic Tails for Human Balance Augmentation. J. Mech. Robot. 16, 061007 (2024).

9. Eden, J. et al. Principles of human movement augmentation and the challenges in making it a reality. Nat. Commun. 13, 1345 (2022).

10. Prattichizzo, D. et al. Human augmentation by wearable supernumerary robotic limbs: review and perspectives. *Prog*. Biomed. Eng. 3, 042005 (2021).

11. Lefumat, H. Z. et al. To transfer or not to transfer? Kinematics and laterality quotient predict interlimb transfer of motor learning. J. Neurophysiol. 114, 2764–2774 (2015).

12. Paparella, G. et al. Relationship between the interlimb transfer of a visuomotor learning task and interhemispheric inhibition in healthy humans. Cereb. Cortex 33, 7335–7346 (2023).

13. Taylor, J. A., Wojaczynski, G. J. & Ivry, R. B. Trial-by-trial analysis of intermanual transfer during visuomotor adaptation. J. Neurophysiol. 106, 3157–3172 (2011).

14. Krakauer, J. W., Mazzoni, P., Ghazizadeh, A., Ravindran, R. & Shadmehr, R. Generalization of Motor Learning Depends on the History of Prior Action. PLoS Biol. 4, e316 (2006).

15. Dempsey-Jones, H. et al. Transfer of tactile perceptual learning to untrained neighboring fingers reflects natural use relationships. J. Neurophysiol. 115, 1088–1097 (2016).

16. Harrar, V., Spence, C. & Makin, T. R. Topographic generalization of tactile perceptual learning. J. Exp. Psychol. Hum. Percept. Perform. 40, 15–23 (2014).

17. Wing, A. M. Motor control: Mechanisms of motor equivalence in handwriting. Curr. Biol. 10, R245–R248 (2000).

18. Wong, A. L., Haith, A. M. & Krakauer, J. W. Motor Planning. The Neuroscientist 21, 385– 398 (2015).

19. Dowdall, L. et al. Developing a Sensory Representation of an Artificial Body Part. Preprint at 10.1101/2025.06.16.658246 (2025).

20. Fan, Z., Lin, C. & Fu, C. A Gaze Signal Based Control Method for Supernumerary Robotic Limbs. in 2020 3rd International Conference on Control and Robots (ICCR) 107–111 (IEEE, Tokyo, Japan, 2020). doi:10.1109/ICCR51572.2020.9344272.

21. Jing, H. et al. A Mouth and Tongue Interactive Device to Control Wearable Robotic Limbs in Tasks where Human Limbs Are Occupied. Biosensors 14, 213 (2024).

22. Meraz, N. S., Shikida, H. & Hasegawa, Y. Auricularis muscles based control interface for robotic extra thumb. in 2017 *International Symposium on Micro-NanoMechatronics and Human Science (MHS)* 1–3 (IEEE, Nagoya, 2017). doi:10.1109/MHS.2017.8305192.

23. Parietti, F., Asada, H. H. Independent, voluntary control of extra robotic limbs. in 5954– 5961 (IEEE, 2017). doi:10.1109/ICRA.2017.7989686.

24. Russ, J. et al. Muscle control of an extra robotic digit. Preprint at 10.1101/2025.06.18.658087 (2025).

25. Zhang, K., Long, Y. & Luo, X. Review of Supernumerary Robotic Limbs. J. Phys. Conf. Ser. 2456, 012004 (2023).

26. Kumar, N., Kumar, A., Sonane, B. & Mutha, P. K. Interference between competing motor memories developed through learning with different limbs. J. Neurophysiol. 120, 1061– 1073 (2018).

27. Wang, J. & Sainburg, R. L. Limitations in interlimb transfer of visuomotor rotations. Exp. Brain Res. 155, 1–8 (2004).

28. Haith, A. M. & Krakauer, J. W. Model-Based and Model-Free Mechanisms of Human Motor Learning. in Progress in Motor Control (eds. Richardson, M. J., Riley, M. A. & Shockley, K.) vol. 782 1–21 (Springer New York, New York, NY, 2013).

29. Yang, C. S., Cowan, N. J. & Haith, A. M. De novo learning versus adaptation of continuous control in a manual tracking task. eLife 10, (2021).

30. Raviv, L., Lupyan, G. & Green, S. C. How variability shapes learning and generalization. Trends Cogn. Sci. 26, 462–483 (2022).

31. Seitz, A. R. Perceptual learning. Curr. Biol. 27, R631–R636 (2017).

32. Wulf, G., Shea, C. & Lewthwaite, R. Motor skill learning and performance: a review of influential factors: Motor skill learning and performance. Med. Educ. 44, 75–84 (2010).

33. Chou, S. et al. The role of the great toe in balance performance. J. Orthop. Res. 27, 549– 554 (2009).

34. Morehead, J. R., Taylor, J. A., Parvin, D. E. & Ivry, R. B. Characteristics of Implicit Sensorimotor Adaptation Revealed by Task-irrelevant Clamped Feedback. J. Cogn. Neurosci. 29, 1061–1074 (2017).

35. Dominijanni, G. et al. The neural resource allocation problem when enhancing human bodies with extra robotic limbs. *Nat*. Mach. Intell. 3, 850–860 (2021).

36. Makin, T. R., De Vignemont, F. & Faisal, A. A. Neurocognitive barriers to the embodiment of technology. *Nat*. Biomed. Eng. 1, 0014 (2017).

37. Masters, R. & Maxwell, J. The theory of reinvestment. Int. Rev. Sport Exerc. Psychol. 1, 160–183 (2008).

38. Schone, H. R. et al. Biomimetic versus arbitrary motor control strategies for bionic hand skill learning. *Nat*. Hum. Behav. 8, 1108–1123 (2024).

39. Ciullo, A. S. et al. A Novel Soft Robotic Supernumerary Hand for Severely Affected Stroke Patients. IEEE Trans. Neural Syst. Rehabil. Eng. 28, 1168–1177 (2020).

40. Hussain, I., Spagnoletti, G., Salvietti, G. & Prattichizzo, D. Toward wearable supernumerary robotic fingers to compensate missing grasping abilities in hemiparetic upper limb. Int. J. Robot. Res. 36, 1414–1436 (2017).

41. Hwang, S., Min, K.-C. & Song, C.-S. Assistive technology on upper extremity function for stroke patients: A systematic review with meta-analysis. J. Hand Ther. 37, 507–519 (2024).

42. Salvietti, G. et al. Compensating Hand Function in Chronic Stroke Patients Through the Robotic Sixth Finger. IEEE Trans. Neural Syst. Rehabil. Eng. 25, 142–150 (2017).

43. Collinger, J. L. et al. Functional priorities, assistive technology, and brain-computer interfaces after spinal cord injury. J. Rehabil. Res. Dev. 50, 145 (2013).

44. Palacios, J. et al. Grasp Force Assistance via Throttle-Based Wrist Angle Control on a Robotic Hand Orthosis for C6-C7 Spinal Cord Injury. IEEE Trans. Med. Robot. Bionics 7, 149–155 (2025).

45. Morioka, H. et al. Robot-assisted training using hybrid assistive limb ameliorates gait ability in patients with amyotrophic lateral sclerosis. J. Clin. Neurosci. 99, 158–163 (2022).

46. Hrga, I. Wearable Technologies: Between Fashion, Art, Performance, and Science (Fiction). Tekstilec 62, 124–136 (2019).

47. Clode, D. et al. Evaluating initial usability of a hand augmentation device across a large and diverse sample. *Sci*. Robot. 9 (2024).

48. Huang, H.-J. & Mercer, V. S. Dual-Task Methodology: Applications in Studies of Cognitive and Motor Performance in Adults and Children: Pediatr. Phys. Ther. 13, 133–140 (2001).

49. Witteveen, H. J. B., De Rond, L., Rietman, J. S. & Veltink, P. H. Hand-opening feedback for myoelectric forearm prostheses: Performance in virtual grasping tasks influenced by different levels of distraction. J. Rehabil. Res. Dev. 49, 1517 (2012).

50. Wetzels, R. et al. Statistical Evidence in Experimental Psychology: An Empirical Comparison Using 855 *t* Tests. Perspect. Psychol. Sci. 6, 291–298 (2011).

